# Cohort-mean measured macromolecules lead to more robust linear-combination modeling than parameterized and subject-specific ones

**DOI:** 10.1101/2022.07.07.499181

**Authors:** Helge J. Zöllner, Christopher W. Davies-Jenkins, Saipavitra Murali-Manohar, Tao Gong, Steve C. N. Hui, Yulu Song, Weibo Chen, Guangbin Wang, Richard A. E. Edden, Georg Oeltzschner

## Abstract

Expert consensus recommends linear-combination modeling (LCM) of ^1^H MR spectra with sequence-specific simulated metabolite basis function and experimentally derived macromolecular (MM) basis functions. Measured MM basis functions have been derived from metabolite-nulled spectra averaged across a small cohort. The use of subject-specific instead of cohort-averaged measured MM basis functions has not been studied. Furthermore, measured MM basis functions are not widely available to non-expert users, who commonly rely on parameterized MM signals internally simulated by LCM software. To investigate the impact of the choice of MM modeling, this study, therefore, compares metabolite level estimates between different MM modeling strategies (cohort-mean measured; subject-specific measured; parameterized) in a lifespan cohort and characterizes its impact on metabolite-age associations.

100 conventional (TE = 30 ms) and metabolite-nulled (TI = 650 ms) PRESS datasets, acquired from the medial parietal lobe in a lifespan cohort (20-70 years of age), were analyzed in Osprey. Short-TE spectra were modeled in Osprey using six different strategies to consider the macromolecular baseline. Fully tissue- and relaxation-corrected metabolite levels were compared between MM strategies. Model performance was evaluated by model residuals, the Akaike information criterion (AIC), and the impact on metabolite-age associations.

The choice of MM strategy had a significant impact on the mean metabolite level estimates and no major impact on variance. Correlation analysis revealed moderate-to-strong agreement between different MM strategies (r > 0.6). The lowest relative model residuals and AIC values were found for the cohort-mean measured MM. Metabolite-age associations were consistently found for two major singlet signals (tCr, tCho) for all MM strategies, however, findings for highly J-coupled metabolites it was depended on the MM strategy. A variance partition analysis indicated that up to 44% of the total variance was related to the choice of MM strategy. Additionally, the variance partition analysis reproduced the metabolite-age association for tCr and tCho found in the simpler correlation analysis.

In summary, the inclusion of a single high-SNR MM basis function (cohort-mean) leads to more robust metabolite estimation (lower model residuals and AIC values) compared to MM strategies with more degrees of freedom (Gaussian parametrization) or subject-specific MM information. Integration of multiple LCM analyses into a single statistical model potentially improves the robustness in the detection of underlying effects (e.g. metabolite vs age), reduces algorithm-based bias, and estimates algorithm-related variance.

## Introduction

Single-voxel proton MRS allows in-vivo research studies of brain metabolism^1,2^. At clinical field strength (≤3T), spectral modeling is hampered by poor spectral resolution as, particularly for short-echo-time (short-TE) spectra, metabolite signals overlap with a broad background consisting of fast-decaying macromolecule and lipid signals. Linear-combination modeling (LCM) is the consensus-recommended quantification method for short-TE spectra. LCM algorithms estimate spectra as a linear combination of metabolite and macromolecular (MM) basis function, maximizing use of prior knowledge to constrain the model solution^3^.

Recent consensus on the modeling of short-TE spectra emphasizes that the MM basis functions included in the LCM must be either accurately parameterized by taking T_1_ and T_2_ evolution of the MMs into account, or directly measured with matching acquisition parameters ^3,4^. However, acquiring MM spectra is time-consuming and technically challenging especially since stock sequences typically lack pre-inversion modules, and protocols for acquiring metabolite-nulled spectra are not widely available. This results in two common practices in the field, either: 1) MM spectra are measured in a small number of subjects and then averaged into a cohort-mean MM basis function to be included in the analysis of the whole cohort and subsequent studies^4^ to minimize protocol duration; or 2) MM spectra are modeled using a default empirical parameterization as a series of Gaussians ^5^ as described in the most widely used LCM algorithm LCModel^6,7^. The latter approach is more commonly taken, as it does not require any additional time-consuming data acquisition or specialized processing, although it is not clear how adequate such parametrizations are.

Interestingly, it has not been investigated whether including subject-specific measured MMs (mMM) in the LCM has advantages over cohort-mean mMMs, possibly because prohibitively long acquisition times have disincentivized such a study, and current MRS analysis software does not provide easily accessible workflows to do so. A recent large-scale lifespan cohort^8,9^ now provides sufficient publicly available data to compare the two strategies. This dataset further allows investigation of other important issues of MM modeling in short-TE spectra to be investigated:

1. <u>Parameterized vs. measured MM:</u> Only a few low-n studies have systematically compared metabolite estimation between parameterized MM (pMM) and mMM modeling, or studied updated MM parameterizations^10–12^.
2. Other studies focused on individual differences between measured MM spectra, but did not investigate the impact on the metabolite estimation^8,13^.
3. LCM algorithms apply modeling parameters to the basis functions, including a single Gaussian linebroadening parameter to account for B_0_ inhomogeneities in the MRS volume^6,14,15^. This practice becomes problematic when simulated metabolite basis functions are combined with a measured MM background, since the latter already has invivo linewidth. It might therefore be more appropriate to use <u>two separate</u> Gaussian linebroadening parameters (one for metabolites, one for MM).

The overarching goal of this study was to systematically investigate different strategies for the inclusion of MM signals for linear-combination modeling of short-TE PRESS spectra, most notably the use of single-subject MM data. We sought to substantiate expert-consensus recommendations on MM handling^3,4^ and provide readily usable workflows for all strategies in our analysis pipeline ‘Osprey’^14^. To this end, different MM modeling strategies were implemented and applied to a large lifespan dataset from healthy volunteers^8,9^. Macromolecular modeling strategies include cohort-mean and subject-specific measured macromolecules, the most commonly used Gaussian parametrizations, and the use of a single or two separate Gaussian linebroadening parameters. In the absence of a ‘gold standard’, the performance of each strategy was evaluated by comparing descriptive statistics of metabolite estimates, Akaike information criteria (AIC), model residuals, and variance partition analysis. Finally, we investigated whether the choice of MM model had an influence on the observation of metabolite-age associations.

## Methods

### Study participants & data acquisition

In this study, a cohort of 102 healthy volunteers was recruited with equal numbers of male and female participants in each decade of life—the 20s to 60s—following local IRB approval (Shandong Hospital, China, Provincial Hospital). Exclusion criteria included a history of neurological or psychiatric illness, and contraindications for MRI. Details of the acquisition protocol can be found in published studies of the same cohort^8,9^. Briefly, all participants were scanned on a Philips 3T MRI scanner (Ingenia CX, Philips Healthcare, The Netherlands) and the protocol included a T_1_-weighted MP-RAGE scan with 1 mm^3^ isotropic resolution for voxel positioning and tissue segmentation. Short-TE metabolite data were acquired using PRESS localization^16^ (TR/TE 2000/30 ms, 30×26×26mm^3^, 96 transients, 2048 datapoints, 2000 Hz sampling bandwidth, VAPOR water suppression^17^) in the posterior cingulate cortex (PCC). Subcutaneous lipid adjacent to the voxel was suppressed with a slice-selective saturation pulse (20 mm thickness). Additionally, metabolite-nulled MM data were acquired in the same voxel with an inversion time of 600 ms using an adiabatic hyperbolic-secant inversion pulse with a full-width half-maximum (FWHM) of 698 Hz applied at 1.9 ppm to minimize impact on the water signal, which was suppressed using Philips CHESS water suppression^18^. Water reference spectra were acquired for each participant without water suppression or pre-inversion.

### Data analysis

#### Pre-processing

Data were analyzed in MATLAB (MATLAB R2022a, The MathWorks Inc., Natick, Massachusetts, United States) using Osprey (v.2.2.0), an end-to-end open-source MRS analysis toolbox. Pre-processing on the scanner included coil-combination, averaging, and eddy-current-correction using the water reference scan^19^. Water reference data were eddy-current-corrected in Osprey (since they are exported from the scanner without this correction). Residual water signal in the metabolite and metabolite-nulled spectra was removed with a Hankel singular decomposition (HSVD) filter^20^. As previously described^8^, imperfectly-nulled metabolite signals in MM spectra were removed with a linear-combination modeling procedure using simulated metabolite signals (NAA, Cr, CrCH_2_, Cho, Glu) and a highly flexible spline (0.1 ppm knot spacing) to model the MM signals in the metabolite-nulled spectrum between 0.5 and 4.2 ppm. The fitted metabolite signals were subsequently subtracted from the spectra to yield ‘clean’ MM spectra. Finally, the clean MM spectra were modeled with a flexible spline model (again 0.1 ppm knot spacing) to generate a noise-free MM basis function for subsequent LCM of the metabolite spectra (**Supplementary Material 1**). Data quality metrics (SNR and linewidth of the NAA singlet) were calculated for the processed metabolite spectra. SNR was defined as the ratio of the NAA singlet peak height and the standard deviation of the frequency-domain spectrum between -2 and 0 ppm. LW was defined as the average of the simple FWHM and the FWHM of a Lorentzian peak model of the NAA singlet.

#### Short-TE linear-combination modeling strategies

Metabolite data were modeled using the Osprey LCM algorithm, using a basis set of 18 metabolite basis functions generated from a fully-localized 2D density-matrix simulation of a 101 × 101 spatial grid (field of view 50% larger than voxel) implemented in the cloud-based MATLAB toolbox ‘MRSCloud’^21^ derived from FID-A^22^. The following basis functions were included: ascorbate, aspartate, creatine, negative creatine methylene (-CrCH_2_, to account for water suppression/relaxation), GABA, glycerophosphocholine (GPC), glutathione (GSH), Gln, Glu, mI, Lac, NAA, NAAG, phosphocholine, phosphocreatine, phosphoethanolamine (PE), Scyllo (sI), and taurine (**Figure 1**). Detailed descriptions of the frequency-domain LCM algorithm and parameter settings can be found elsewhere^5,14^. Briefly, metabolite spectra were analyzed between 0.5 and 4 ppm with a spline baseline knot spacing of 0.4 ppm. The model frequency range was different from the MM spectra to reduce possible confounds from the different water suppression techniques (CHESS for MM spectra; VAPOR for metabolite spectra)^8^ and allow direct comparison to previous studies^5,23^. While the highly flexible (0.1 ppm knot spacing) spline model needs to fully capture the broad MM signals in the metabolite-nulled spectra, the 0.4-ppm knot spacing setting was chosen as the spline baseline is only required to account for low-frequency background fluctuations in the metabolite spectra. This setting for LCM of short-TE MRS in Osprey has previously led to good agreement with other LCM algorithms^5^. Model parameters of Osprey’s frequency-domain LCM algorithm include amplitude estimates for the metabolite basis functions, frequency shifts, zero/first order phase correction, Gaussian and Lorentzian linebroadening, and cubic spline baseline coefficients. Water reference data were modeled with a simulated water basis function and a 6-parameter model (amplitude, zero- and first-order phase, Gaussian and Lorentzian linebroadening, and frequency shift).

**Figure 1.**
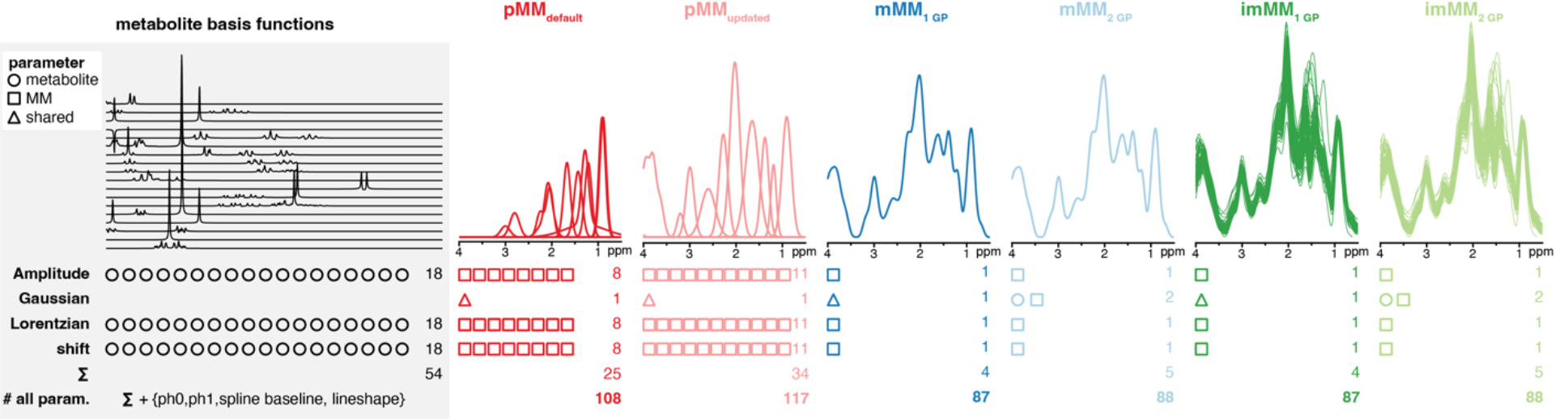
Strategies to include macromolecules in the LCM of short-TE MRS, including commonly used default (pMM_default_ in bold red), updated Gaussian parameterization (pMM_updated_ in light red) based on measured MM spectra, single measured MM based on the cohort mean (mMM_1 GP_ and mMM_2 GP_ in bold and light blue, respectively), and individual measured MM (imMM_1 GP_ and imMM_2 GP_ in bold and light green, respectively). Inclusion of a single Gaussian model parameter for metabolites is marked “1 GP” and models using separate parameters are marked “2 GP”. The number of model parameters, and indicators whether those are shared between metabolite and MM basis *functions* (triangle) or separate (circle and square respectively for metabolite and MM) are shown below.

The centerpiece of this study is the comparison of different implementations to account for MMs in short-TE LCM (**Figure 1**). In general, each MM basis function has its own model parameters for the amplitude, frequency shift, Lorentzian linebroadening. The six strategies can be grouped into ‘parameterized MM’ (pMM) and ’measured MM’ (mMM) strategies, and the mMM strategies have been implemented with one shared or two separate Gaussian linebroadening parameters for metabolites and MM:

- The default parametrization **pMM_default_** model (**bold red**) contains 8 Gaussian basis functions to parameterize the MMs and lipids as previously described^5^. The parameterization is derived from the LCModel manual^7^ without the definition of expectation values for the Lorentzian linebroadening and the frequency shift during the optimization and a simplified Lipid model at 1.3 ppm. Detailed amplitudes, frequency shifts, linewidth, and soft constraints for the LCM can be found in **Supplementary Material 2**. This parametrization is still the most widely used implementation and its performance has previously been described for a large multi-site dataset^5^.
- The updated parametrization **pMM_updated_** model (**light red**) incorporates a total of 11 MM basis functions which are generated by averaging the spline model of all metabolite-nulled spectra and modeling that as a linear combination of 14 Gaussian peaks – between 0.5 and 4.2 ppm - with the initial frequency shifts, as previously described^8^. This MM strategy does not include additional lipid peaks, as the averaged spline model did not allow for the separation of the broad peaks at 0.9 and 1.3 ppm. The Gaussian basis functions between 3.7 and 4.0 ppm overlapped substantially and were therefore summed to generate a single ‘MM_3.7-4.0_’ basis function, an approach described previously^24^. Again, detailed parametrization can be found in **Supplementary Material 2**.
- The cohort-mean measured **mMM_1 GP_** model (**dark blue**) has a single MM basis function generated by averaging the noise-free metabolite-nulled spectra from all participants. The same shared Gaussian lineshape parameter is applied to the metabolite and the MM basis function (therefore named mMM_1 GP_). This approach has the lowest number of model parameters and benefits from high signal-to-noise by using the averaged MM spectra. The creation of this across-age-cohort-mean MM basis function is only appropriate as no significant changes in the MM signals were observed as a function of age^8^. However, it enforces fixed ratios between the MM signals and is unable to capture any subject-specific variation of the MM signals. This approach has previously been employed in the same cohort^9^ and closely corresponds to the MRS data analysis consensus recommendations^3,4^.
- The cohort-mean measured **mMM_2 GP_** model (**light blue**) is identical mMM_1 GP_, except that two Gaussian lineshape parameters are included to separately model additional linewidth applied to metabolite basis functions and the MM basis function.
- The individual subject-specific measured **imMM_1 GP_** (**dark green**) uses the noise-free metabolite-nulled spectrum from each individual subject, instead of the cohort-averaged MM basis function. This strategy incorporates subject-specific MM variation.
- The subject-specific measured **imMM_2 GP_** model (**light green**) implements the same subject-specific MM basis function with separate Gaussian lineshape parameters applied to metabolite and MM basis functions.

All models are implemented in Osprey v2.2.0 and freely available on GitHub^25^. A summary page following the minimum reporting standards in MRS consensus paper^26^ can be found in **Supplementary Material 3**.

### Quantification, visualization, and statistics

#### Quantification

Water-referenced metabolite concentrations were calculated using tissue-specific water visibility and relaxation based on literature values^27,28^ for each segmented tissue fraction of the voxel derived from the SPM12-based co-registration and segmentation of the T_1_-weighted anatomical image and the MRS voxel^29^.

#### Visualization

The model performance and spectral characteristics arising from the systematic differences of each modeling strategy were visually assessed through the mean spectra, mean fit model, mean total MM (tMM) models, mean spline baseline, mean total background (tMM + spline baseline), and mean residuals (means taken across all datasets).

Metabolite estimates are visualized as raincloud plots^30^ with median, 25^th^ and 75^th^ percentile ranges, individual estimates, smoothed distributions, mean and standard deviation to identify systematic differences between the modeling strategies. The corresponding coefficients of variation (CV) are reported as bar plots. Correlations between metabolite estimates derived using the different approaches were reported as bar plots representing Pearson’s correlation coefficient, *r*.

Statistical differences between all pairs of correlation coefficients were evaluated using a modified Fisher’s Z procedure^31^ as implemented in the cocor package^32^ and corrected for multiple comparisons using Bonferroni correction. In addition, a correlation matrix comparing the correlation between the participant age and the metabolite/MM estimates for all different approaches was created. The correlation matrix includes Pearson’s correlation coefficients, *r*. For significant associations after applying Bonferroni correction (treating each MM strategy as a separate analysis with a correction factor of 15), the directionality of the correlation is encoded as a color-coded ellipsoid, with higher associations represented by a narrower ellipsoid, i.e. a circle indicates there is no association (r = 0) and a line indicates perfect association (r = 1), additionally emphasized by the size of the ellipsoid (small ellipsoid ≡ small r). The p values were encoded by the linewidth of the ellipsoid (the smaller the p value, the stronger the line). Metabolite-age associations were further tested for stability using a variable selection difference (VSD) approach by sampling random subsets of the data as implemented in the glmvsd package^33^. The VSD was calculated for all metabolites and MM strategies removing 5%, 20% and 50% of the subjects with 5000 simulations each. VSD scores close to 0 indicate that no additional predictor is needed as age is a stable predictor for the metabolite concentration even in randomly sampled subsets of the data. VSD scores closer to 1 indicate that age is a poor predictor for the metabolite concentration and additional predictors are needed. All plots and statistics were generated in R^34^ (Version 4.1.3) in RStudio (Version 2022.02.01, Build 461, RStudio Inc.) using SpecVis^5^, an open-source package to visualize linear-combination modeling results with the ggplot2 package^35^. All scripts and results are publicly available on the Open Science Framework^36^.

#### Statistics

Differences in the mean and the variance of the metabolite level estimates were assessed between pMM_default_ and all other MM strategies. The statistical tests were set up as paired without any further inference. Fligner-Killeen’s test with a post-hoc pair-wise Bonferroni corrected Fligner-Killeen’s test was employed to test differences in metabolite estimate variance. An ANOVA, or Welch’s ANOVA (depending on whether variances were different or not) was employed to test differences in the means. Post-hoc analysis was performed with a Bonferroni corrected paired t-test using non-equal or equal variances, respectively.

#### Variance partition analysis with linear mixed-effect model

Linear mixed-effect (LME) models were set up as a repeated-measure analysis to determine variance partition coefficients to assess the contribution of MM strategy-, age-, sex- and participant-related contributions to the total variance of metabolite estimates as described previously^5,23,37,38^. Age and sex were defined as fixed effects while MM approach and participant were defined as random effects. The probability of observing the test statistic was evaluated against the null hypothesis, which was simulated by performing parametric bootstrapping (10,000 simulations)^39^.

The resulting p-values were corrected for multiple comparison using the Benjamini-Yekutieli approach^40^ which is less conservative than a Bonferroni correction, taking into account the dependency between the metabolite level estimates.

### Model evaluation criteria

The performance of each modeling approach was evaluated using several metrics including the impact on the metabolite estimates, metabolite-age associations, as well as several quality metrics:

1. Visual inspection for characteristic spectral features included: Agreement between mean models, structured residual, flexibility and agreement of the spline baseline, tMM, and background (spline baseline + tMM).
2. Consistency of metabolite estimates: between-subject coefficients of variation (CV = SD/mean) for all metabolite estimates were calculated for each modeling approach. Between-subject CV was interpreted as a measure of modeling performance, assuming that increased CVs are mainly introduced by variability in the modeling and do not reflect biological meaningful variance of metabolite estimates.
3. Agreement in the metabolite estimation between the different modeling approaches were assessed via a correlation analysis.
4. Amplitude fit quality: The difference between the maximum and minimum of the residual was determined, and then normalized by the noise level^2^. This was done over the entire modeling range and termed relative residual.
5. AIC: The Akaike information criterion^41^, which takes the number of model parameters into account, is defined as follows:

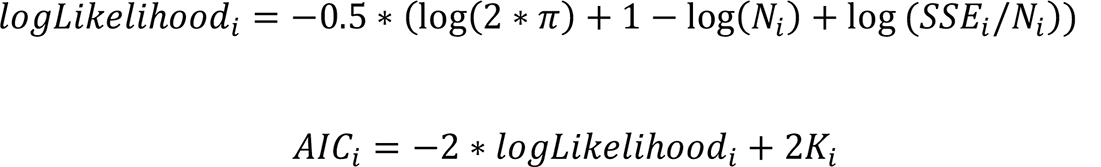 Here, *N_i_* is the number of points in the i^th^ modeling strategy, *SSE_i_* is the sum of squared error (i.e., squared residual) of that strategy, and *K_i_* is the number of free model parameters for that strategy. No further corrections for amplitude soft-constraints were employed. Lower *AIC_i_* values indicate a more appropriate model. Subsequently, ***ΔAIC_i_*** scores were calculated as the difference of *AIC_i_* of modeling strategy *i* and the model with the lowest *AIC_min_*:

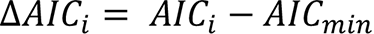
6. Inter-class correlation coefficients (ICC) were calculated between the clean MM spectra and the tMM model from the metabolite spectra LCM, as well as the clean MM spectra and the sum of tMM and spline baseline (spbl) model for all subjects.
7. Age-metabolite and age-MM associations were assessed for all models to identify interactions between differences in the model attributing signal to metabolites, MM, or spline baseline, as well as possible bias in a specific approach.
8. LME models were used to determine variance partitions for MM strategy, age, sex, participant, and residual unexplained variance which allows to make inferences about all effects across all datasets and determine influential factors in the quantitative analysis of large-cohort data.

## Results

All 102 datasets were successfully analyzed. After visual inspection, one subject was removed due to a large visible ethanol signal and a second subject was removed due to unwanted out-of-volume echoes visible in the model residual leading to a final total of 100 datasets. The data quality assessment indicated consistently high spectral quality (SNR_NAA_ = 151 ± 23; LW_NAA_ = 6.7 ± 0.9 Hz) without lipid contamination.

### Visualization of modeling

A summary of the data and modeling results is presented in **Figure 2**. Overall, a strong agreement between the mean of data and model is found for all modeling strategies. However, there is a substantial structured signal in the mean residual at 0.9 ppm for the pMM_updated_ strategy, indicating non-optimal parametrization of the MM_0.93_ peak with a single MM basis function in this frequency region. The mean of the sum of the MM and baseline models agree well across all strategies, indicating that the combination of these two terms achieves comparable modeling of all broad non-metabolite signals, although the pMM strategies exhibit a broader and differently shaped signal in the 2-ppm region compared to the mMM strategies. Of note, there appears to be no systematic difference between the cohort-mean and the subject-specific mMM strategies, or between the strategies using 1 or 2 Gaussian linebroadening parameters. Upon closer inspection, it becomes strikingly obvious that the main differences appear between the pMM and mMM strategies, particularly with regard to how much signal they attribute to the baseline and the MM model, respectively, i.e. all pMM strategies attribute a lot more signal to the baseline model. In contrast, the mMM strategies exhibit a much flatter baseline model, and closely follow the shape of the mean mMM signal (except for a small scaling factor related to the different relaxation experienced by the MM between the metabolite-nulled and the metabolite spectra and the potential impact on the baseline introduced by the different water suppression schemes). Interestingly, the pMM_default_ strategy agrees well with the mMM strategies at 0.9 ppm, but substantially less in the remaining frequency regions. The smaller residual could be related to a more accurate parametrization of the MM signal in this region, as two separate Gaussians (MM_0.91_ & Lip_0.89_) are included at 0.9 ppm for this strategy. The mean spline baseline models of the parameterized strategies have a higher number of more pronounced peaks (1.4, 2.2, 3.0, and 3.7 ppm) compared to the experimentally derived strategies indicating that the LCM algorithm estimates a larger portion of the overall signal as baseline for the former strategies.

**Figure 2.**
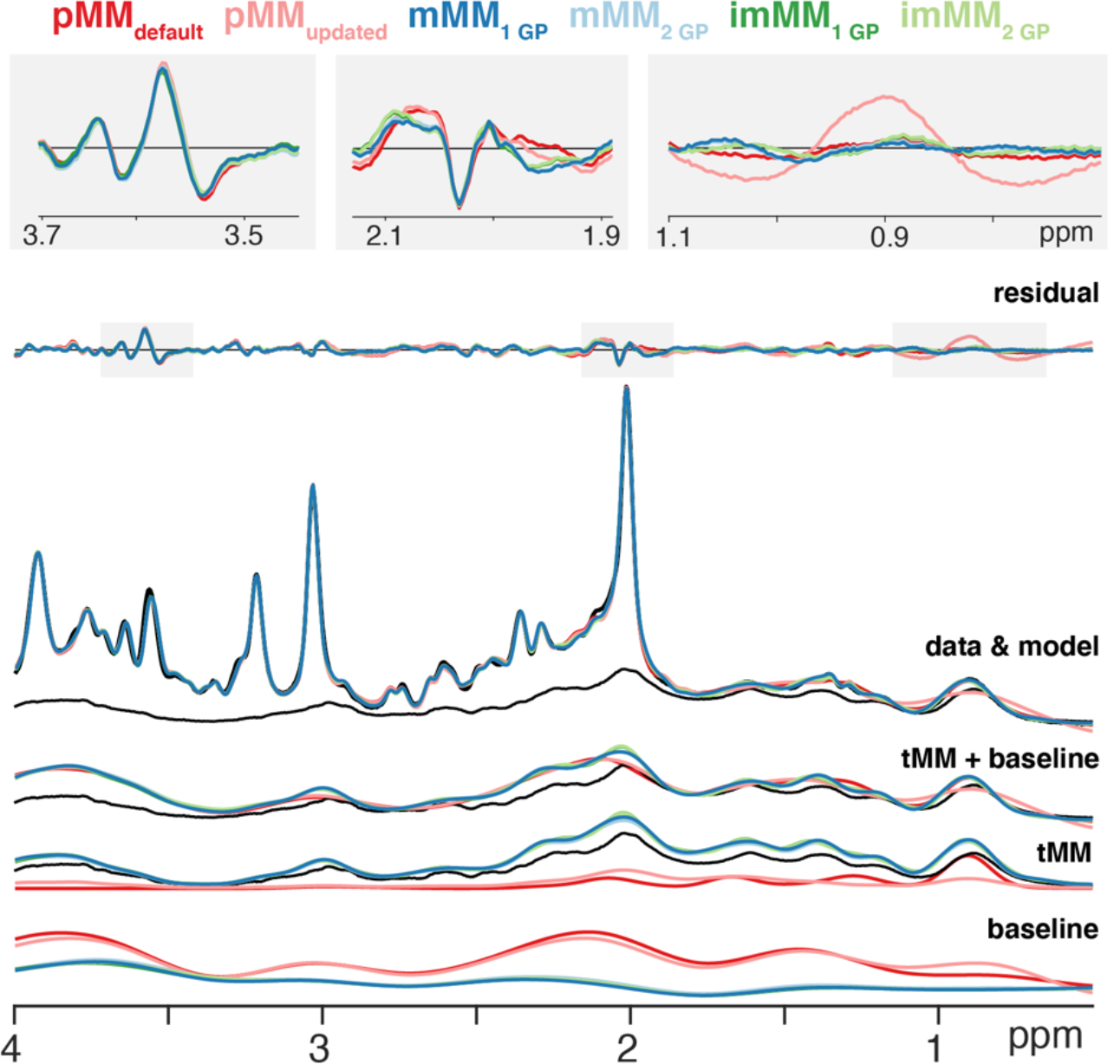
Mean modeling results for all modeling strategies. Data and model agree well for all strategies; however, a substantial structured residual is apparent at 0.9 ppm for the pMM_updated_ strategy. All six models are summarized with the mean results for residual, data (short-TE metabolite and MM), model, total MM model (tMM) + spline baseline model, tMM model, and spline baseline model. In addition, the mean of the ‘clean’ MM spectra is shown in black for reference. Three frequency regions with the largest residuals are magnified in the inlays at the top.

### Metabolite level estimates: Distribution and Correlation

A summary analysis of the metabolite level estimates comparing the ‘default parametrization’ strategy pMM_default_ to all other strategies is shown in **Figure 3** for the major metabolites and in **Figure 4** for all other metabolites. **Table 1** reports numerical summary of the results. Significant differences in mean metabolite level estimates between pMM_default_ and any other strategy were found for almost all metabolites (**Figure 3A, 4A, and 4D**).

**Figure 3.**
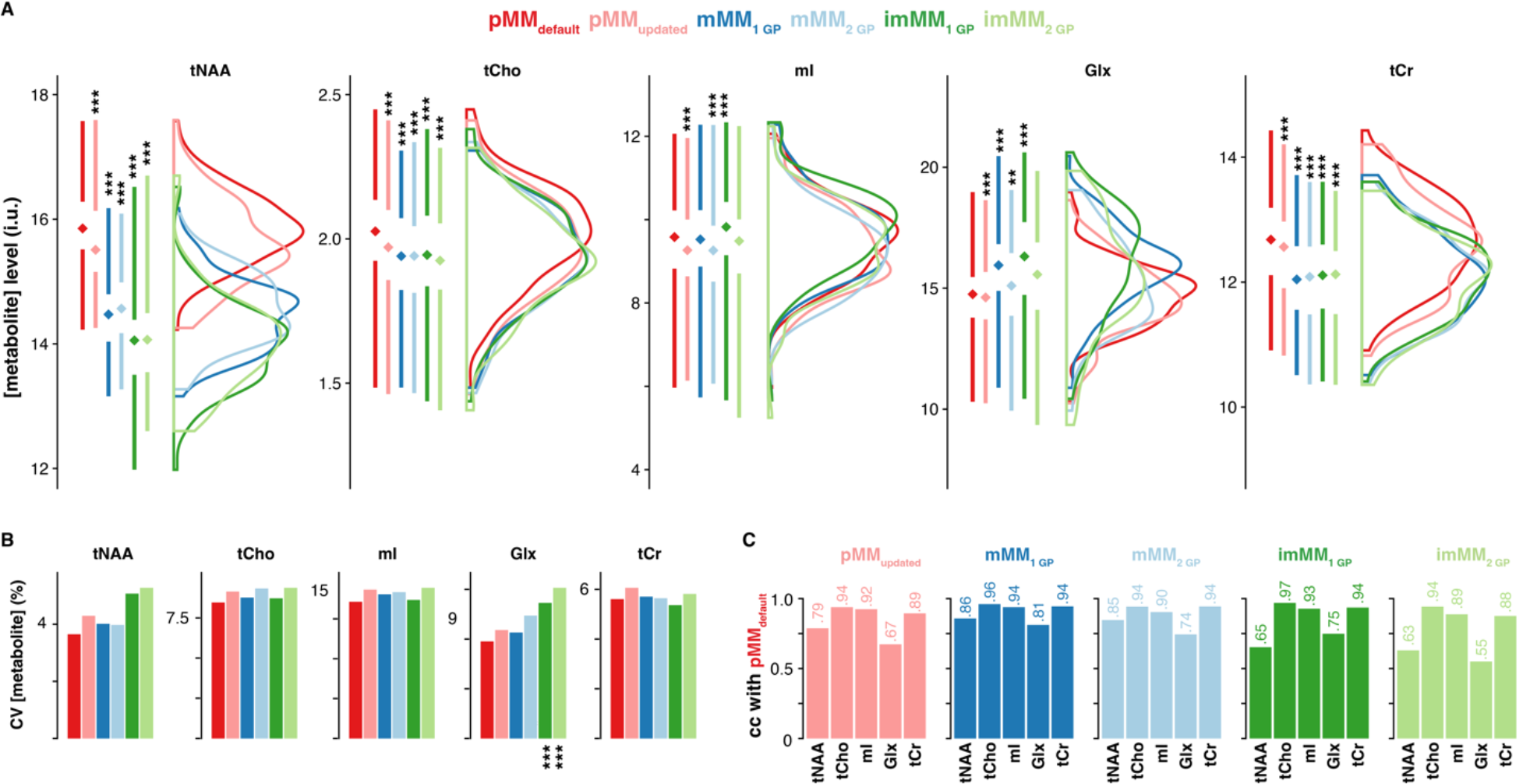
Metabolite level estimates distribution and correlation between MM strategies for the major metabolites. (A) Metabolite estimate distributions major for all MM strategies. For the raincloud plots, box plots with median, interquartile range, and 25^th^/75^th^ percentile, and smoothed distributions are shown. (B) Bar plots of the corresponding CV values. Asterisks indicate significant differences in the mean or variance between the pMM_default_ and any other MM strategy (adjusted p < 0.05 = *, adjusted p < 0.01 = **, adjusted p < 0.001 = ***). (C) Pearson’s correlation coefficient, r, for the correlation analysis between the pMM_default_ and all other MM strategies. All correlations were found to be significant.

**Figure 4.**
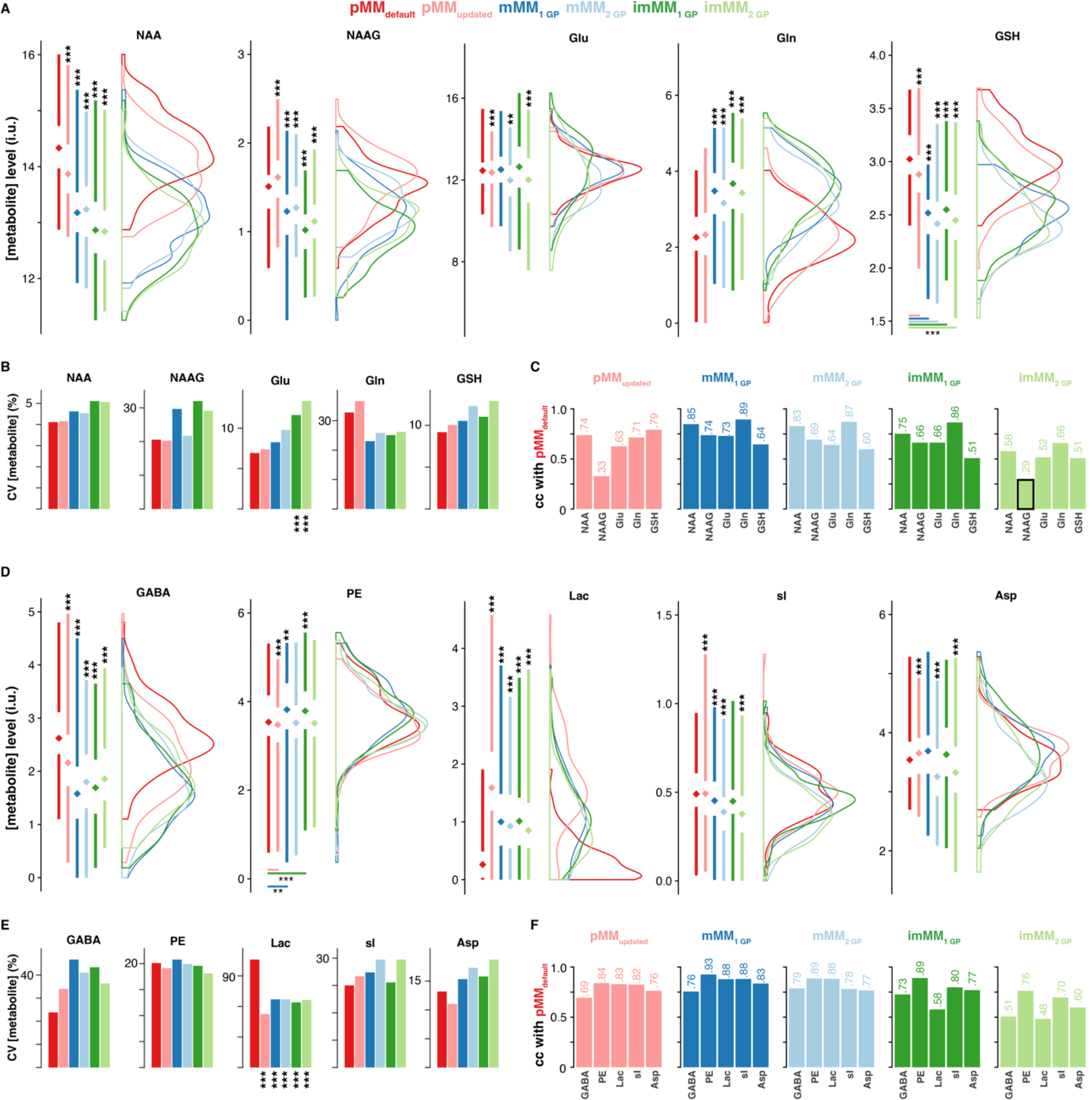
Metabolite level estimates distribution and correlation between MM strategies for all other metabolites. (A) Metabolite estimate distributions major for all MM strategies. For the raincloud plots, box plots with median, interquartile range, and 25^th^/75^th^ percentile, and smoothed distributions are shown. (B) Bar plots of the corresponding CV values. Asterisks indicate significant differences in the mean or variance between the pMM_default_ and any other MM strategy (adjusted p < 0.05 = *, adjusted p < 0.01 = **, adjusted p < 0.001 = ***). (C) Pearson’s correlation coefficient, r, for the correlation analysis between the pMM_default_ and all other MM strategies. All correlations were found to be significant .

**Table 1.**
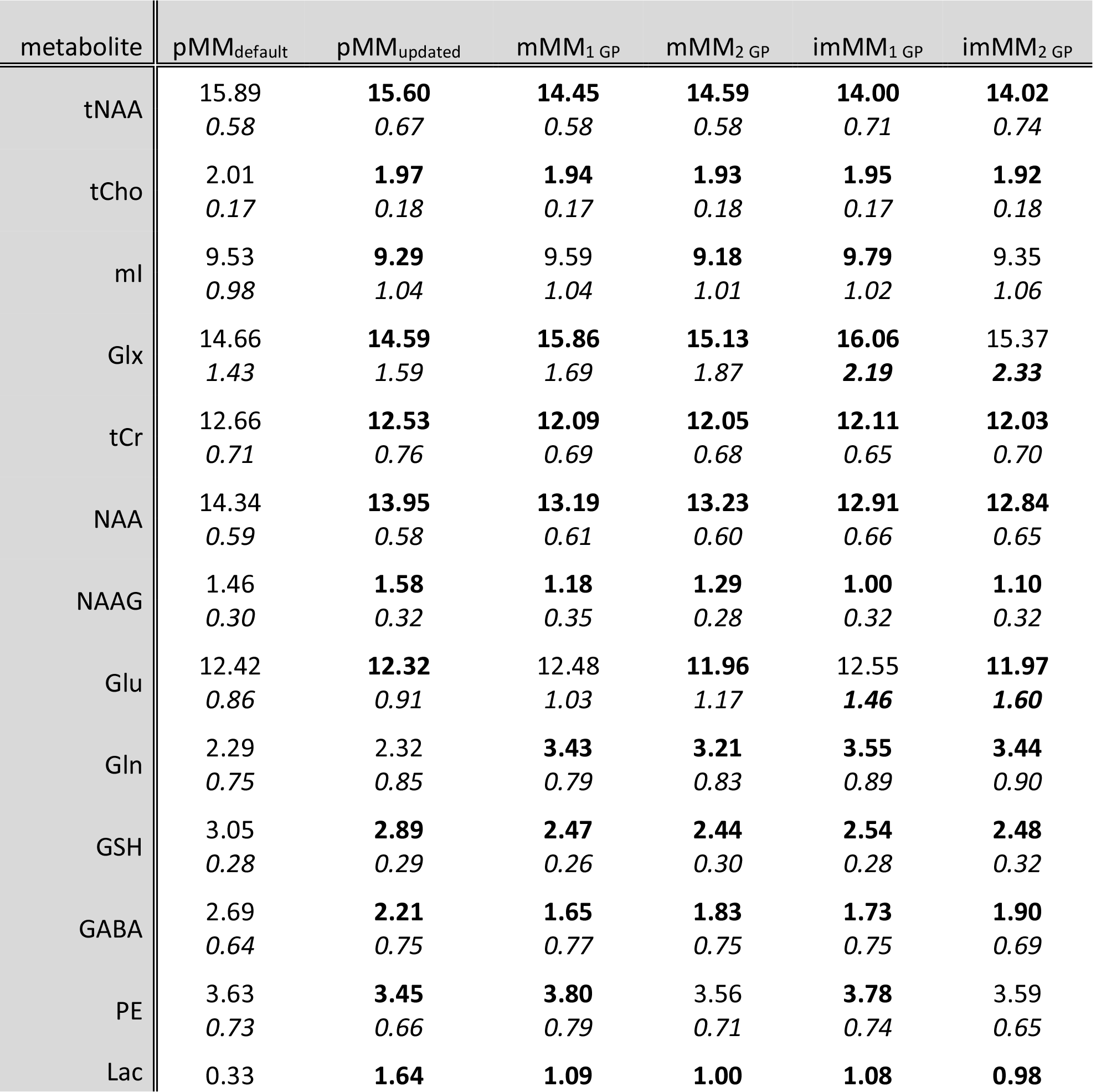
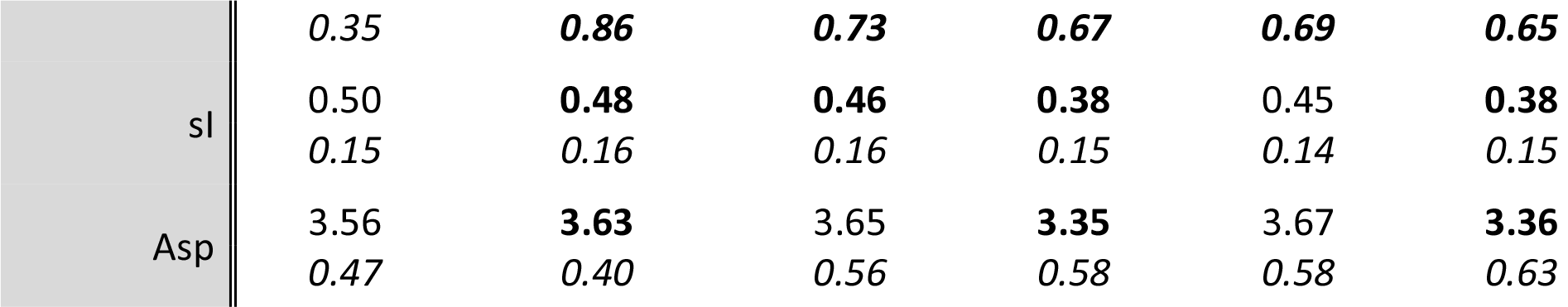
Summary of the descriptive statistics of the metabolite level estimates. In each case, the upper number is the mean metabolite concentration, and the lower italic number is the standard deviation of the metabolite concentration across all subjects. Significant differences in mean or variance between a specific MM strategy and the pMM_default_ strategy are marked in bold.

Most notably, these include reductions in tNAA, tCr (**Figure 3A**), separate NAA and NAAG, and GSH (**Figure 4A**). A significant increase in the metabolite levels compared to the default parametrization strategy was found for Glx which is mainly driven by changes in the Gln estimates. No overall trend for the variance of the metabolite level estimates was observed. A significant increase in variance was found for three metabolites when using subject-specific measured MM spectra, namely Glx (imMM_1 GP_, imMM_2 GP_), Glu (imMM_1 GP_,imMM_2 GP_), and lactate (pMM_updated_, mMM_1 GP_, mMM_2 GP,_ imMM_1 GP_, imMM_2 GP_). This is potentially a result of the low SNR of individual MM spectra causing unwanted variance during the removal of residual metabolite signal during the MM clean-up. Lactate metabolite level estimates are substantially reduced for the pMM_default_ strategy and exhibit significantly higher variance compared to all other MM strategies (**Figure 4D** and **E**).

In general, metabolite estimates correlated strongly (r > 0.6) for almost all metabolites between pMM_default_ and all other strategies (**Figure 3C**. **4C, and 4F**). The associations were particularly strong (r > 0.85) across all MM strategies for three (tCho, tCr, and mI) of the major metabolites either characterized by a strong singlet signal or total absence in the MM spectra. Lower associations (r < 0.8) were found for NAA, NAAG, and tNAA in pMM_updated_, imMM_1 GP_ and imMM_2 GP_. Furthermore, Glx showed the weakest associations of the major metabolites between the MM strategies. Strategies generally agreed less well for low-concentration highly J-coupled metabolites (GSH, and NAAG) since their complex multiplet signals are harder to reliably separate from the broad background. Supporting this notion, weaker between -strategy correlation was also found for J-coupled metabolites whose residual contributions are difficult to reliably and accurately remove from the MM spectra (i.e. Glx. The analysis of the correlation coefficients (**Supplementary Material 4**) emphasizes these findings, indicating strong agreement for tCho, tCr, and mI. Significant differences where mainly found between pMM_default_ and all other strategies, and between imMM_2 GP_ and all other strategies.

The introduction of separate Gaussian linewidth parameters for metabolites and MM resulted in significantly lower concentration estimates for 13 out of 30 metabolites, unchanged concentrations for 13 out of 30 metabolites and significantly higher concentrations for 4 out of 30 metabolites when compared to the MM strategies with a single shared Gaussian linewidth parameter and two separate Gaussian linewidth parameter . Metabolites with strong singlet signals were not affected by this, except for tNAA, driven primarily by changes in the NAAG level estimation. Interestingly, the direction of change when introducing a second Gaussian parameter was similar for both the mMM and imMM strategies. 2-GP strategies did not result in changes in metabolite level variance compared to 1-GP strategies, except for the case of lactate where the additional Gaussian parameter reduced the observed variance of both cohort-mean and individual-measured models (mMM_1 GP_ vs mMM_2 GP_ and imMM_1 GP_ vs imMM_2 GP_).

### Model evaluation criteria

The results of the model evaluation criteria in terms of model residual, AIC, and agreement between short-TE LCM tMM model and individual MM spectra are shown in **Figure 5**. A significant reduction of the model residual was found for all mMM strategies compared to the pMM strategies (**Figure 5A**). This suggests that an accurate representation (mMM and imMM) with fewer of degrees of freedom outperforms generalized representations (pMM) with a higher number of degrees of freedom. The imMM_1 GP_ strategy resulted in the overall lowest AIC, therefore, it was taken as the reference (AIC_min_) for the ΔAIC calculation (**Figure 5B**). However, no significant difference in the AIC was found between the subject-specific imMM_1 GP_ and cohort-mean mMM_1 GP_ strategies, indicating comparable parsimony for both strategies. A significant but small increase in the ΔAIC value was found for the strategies with separate Gaussian linewidth parameters for metabolites and MM (imMM_2 GP_ & mMM_2 GP_) indicating that the possible advantage of one additional linewidth parameter was not justified by sufficiently high improvements of the model residual. Both parameterized strategies resulted in significantly higher ΔAIC values relative to the experimentally derived strategies due to the larger number of MM components and higher degree of freedom without offering lower model residuals than the mMM and imMM strategies.

**Figure 5.**
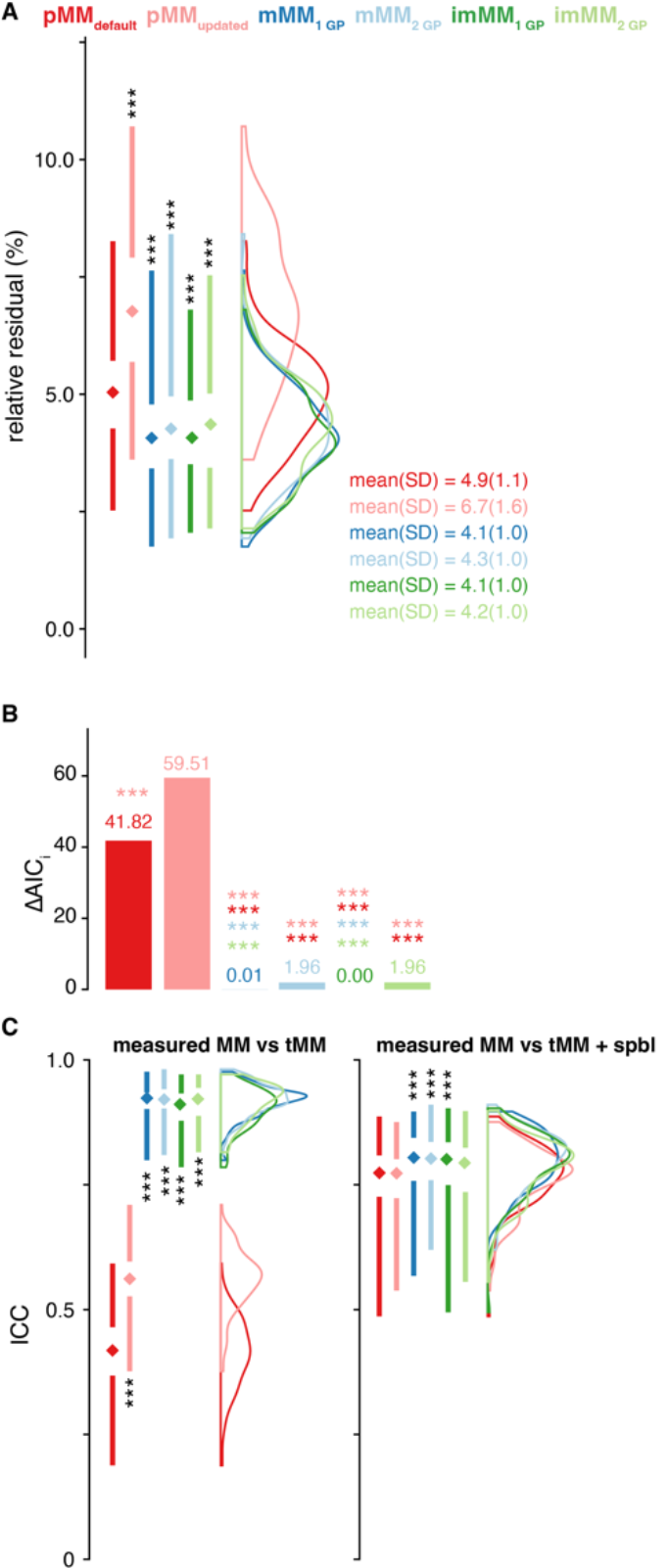
Secondary analysis of residuals, and agreement between short-TE LCM total MM (tMM) model and individual MM spectra. Lower residual amplitudes and AIC scores were found for experimentally derived MM strategies. (A) Relative residual (%) for all MM strategies. (B) Mean ΔAIC values for all MM strategies. Asterisks indicate significant differences (adjusted p < 0.001 = ***) between a specific pair of MM strategies (color-coded). (C) ICC between the individual’s MM data and the short-TE LCM tMM or tMM + spbl model of the same individual. Asterisks indicate significant differences (adjusted p < 0.05 = *, adjusted p < 0.001 = ***).

The ICC analysis (**Figure 5C**) substantiates observations from Figure 2, i.e. poor agreement between the short-TE LCM tMM model estimated by the pMM strategies and the ‘ground-truth’ measured MM data. This indicates that the LCM performs relatively well in separating metabolite and broader background signals, however, it is unable to distinguish between spline baseline and MM signals, especially for the pMM strategies. When the sum of the MM and baseline models is considered, the agreement with the measured MM background improves, but remains significantly poorer for pMM_default_ compared to 3 out of 4 measured MM strategies (mMM_1 GP_, mMM_2 GP_, imMM_1 GP_).

Finally, Gaussian lineshape parameter estimates did not differ between MM strategies. The mean shared Gaussian lineshape width were 8.94 ± 1.35 Hz for pMM_default_, 8.93 ± 1.31 Hz for pMM_updated_, 8.93 ± 1.37 Hz for mMM_1 GP_, and 8.93 ± 1.36 Hz for imMM_1 GP_. The mean separate Gaussian lineshape width for the metabolites were 8.91 ± 1.30 Hz for mMM_2 GP_ and 8.90 ± 1.34 for imMM_2 GP_. The mean separate Gaussian lineshape width for the MM were 5.09 ± 0.07 Hz for mMM_2 GP_ and 5.08 ± 0.08 Hz for iMM_2 GP_ and significantly smaller than for the metabolites. This supports the assumption that less Gaussian line broadening is needed for the experimentally derived MMs; however, this did not affect the metabolite Gaussian parameter.

### Effect on metabolite-age associations

The metabolite-age associations for all MM strategies are summarized in a correlation matrix in **Figure 6**. The two strong positive correlations between metabolite level and age for the two major singlet metabolites tCho (*r* between 0.35 and 0.39) and tCr (*r* between 0.51 and 0.62) are stable across all MM strategies with the strongest association found for the mMM_1 GP_ strategy. Metabolite-age associations are more variable between MM strategies for highly J-coupled metabolites that are affected by the characterization of the MM background for the frequency region > 3.2 ppm (see MM_3.7-4.0_ age association in **Supplementary Material 5**). This is especially apparent for mI (*r* between 0.07 and 0.22), Glx (r between -0.15 and -0.49, affecting Glu and Gln similarly), NAAG (*r* between -0.01 and -0.31), and GSH (*r* between 0.07 and 0.32). It is worth noting that changes in the tCr association may impact the interpretation, if creatine ratios are reported. Additionally, investigating changes in metabolites of interest in aging (such as GSH as a marker of oxidative stress^42,43^) may arrive at different conclusions depending on the MM modeling approach (although, note that quantification of GSH with short-TE LCM at 3T is controversial). Again, as found in **Figure 3B** and **4B**, the separability of metabolite signal and background is driving the differences in the metabolite-age associations, as for example mI appears to have lower variability as for example Glx and GSH. Similar to the changes for > 3.2 ppm frequency region, the variability in the non-significant correlation of age and tNAA may be explained by signal contributions in the 2 ppm frequency region being attributed to the MM or baseline (see MM_2.0_ age association in **Supplementary Material 5**), especially as the MM_2.0_ component is significantly associated with age for the pMM strategies.

**Figure 6.**
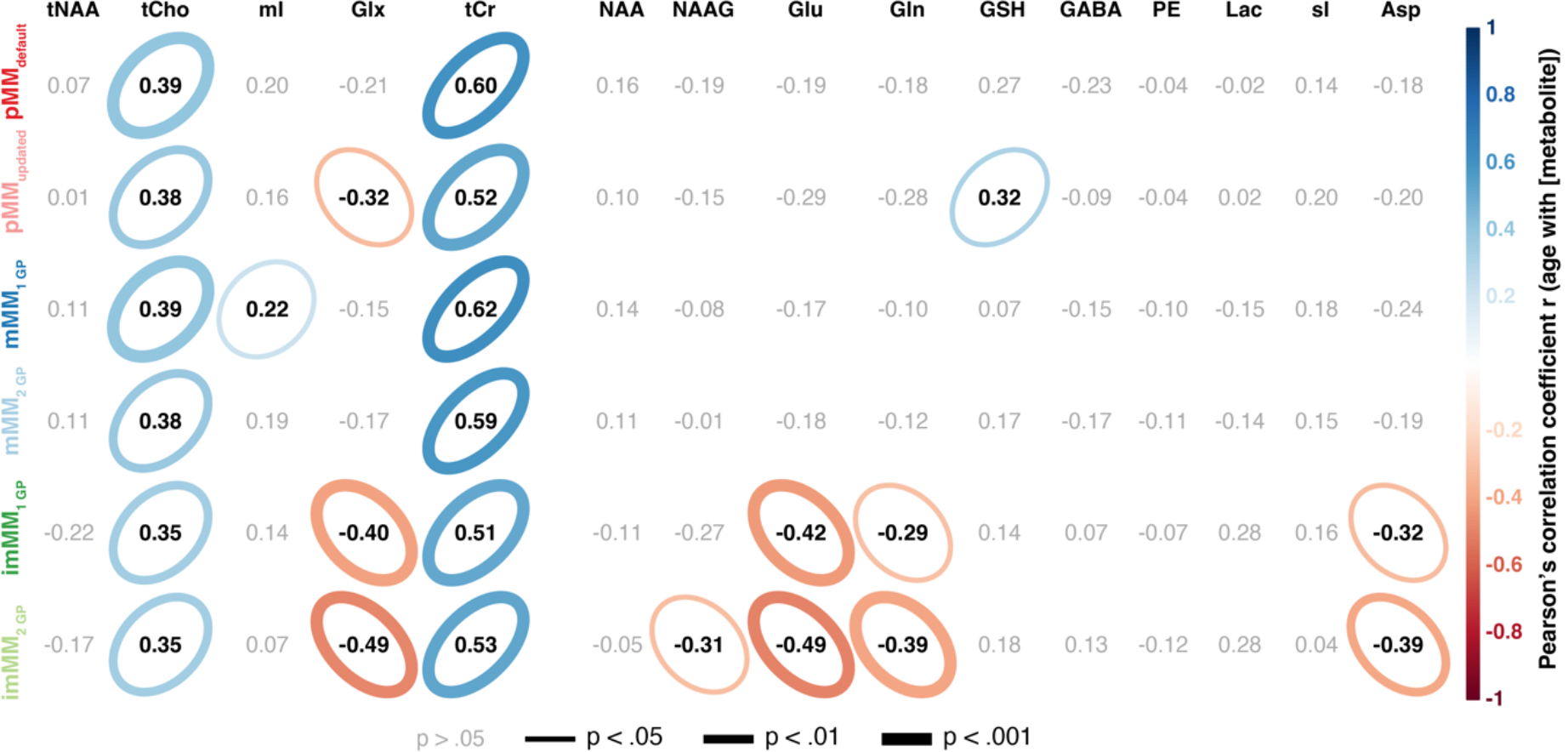
Secondary analysis of metabolite-age associations for all MM strategies. Correlation matrix of Pearson’s correlation coefficient, r, for the metabolite-age association. Results indicate stable metabolite-age associations for the major singlet metabolites, but more variable results for metabolites with stronger J-coupling. Significant correlations are indicated by color-coded ellipsoids with the directionality of r indicated by color and orientation and magnitude of r by the size of the ellipsoid. The p values were encoded by the linewidth of the ellipsoid (the smaller the p value the stronger the line). Bonferroni correction is employed for each MM strategy separately (correction factor of 15 metabolite level estimates).

The stability of the metabolite-age associations are summarized in **Supplementary Material 6.** VSD scores of 0 or close to 0 (Gln for imMM_1 GP_) were found in case of 5% and 20% data removal for all metabolite-age associations that were found to be significant (see Figure 6). Even upon removal of 50% of the data, the VSD remained within acceptable limits (0 for tCr, <0.06 for tCho, and <0.11 for all other) for those associations. The VSD of the remaining associations ranged between .11 and 0.91 indicating lower stability.

The variance partition analysis is summarized in **Figure 7** and **Table 2**. For most metabolites, the largest portion of the variance is explained by the participant level, representing biological variance between subjects. On average ‘participant’ explained 52.3% of variance, MM strategy 19.3%, age 5.2%, and sex 0.95% while 22.3% of variance was not accounted for by these factors. This indicates good responsive of all models to biological variance, significant differences in the metabolite estimates for different MM strategies contributing to overall variance, and no general age or sex ‘pattern’ across all metabolites. Choice of MM strategy explains a significant portion of the variance for all metabolites and is a stronger contributor to the overall variance than ‘participant’ for tNAA (56.3% vs 29.4%), NAA (44.4% vs 38.6%), and GSH (40% vs 31.4%). In addition, 43.7% of the variance of Glu is not explained by any of the included factors. Comparing this analysis, which combines all results in a single analysis, with the simpler correlation analysis separated by MM strategy (Figure 5) confirms some of the metabolite-age associations. For example, tCr and tCho show a strong impact of age, which was also consistently found across all MM strategies in the correlation analysis. In contrast to the correlation analysis, the LME analysis also indicates significant contributions of age effects for Glx, Glu, Gln, and Asp which were found only for a subset of MM strategies in the correlation analysis. Further, no variance contribution from age was found for mI, NAAG, and GSH, which were in turn observed in the correlation analysis.

**Figure 7.**
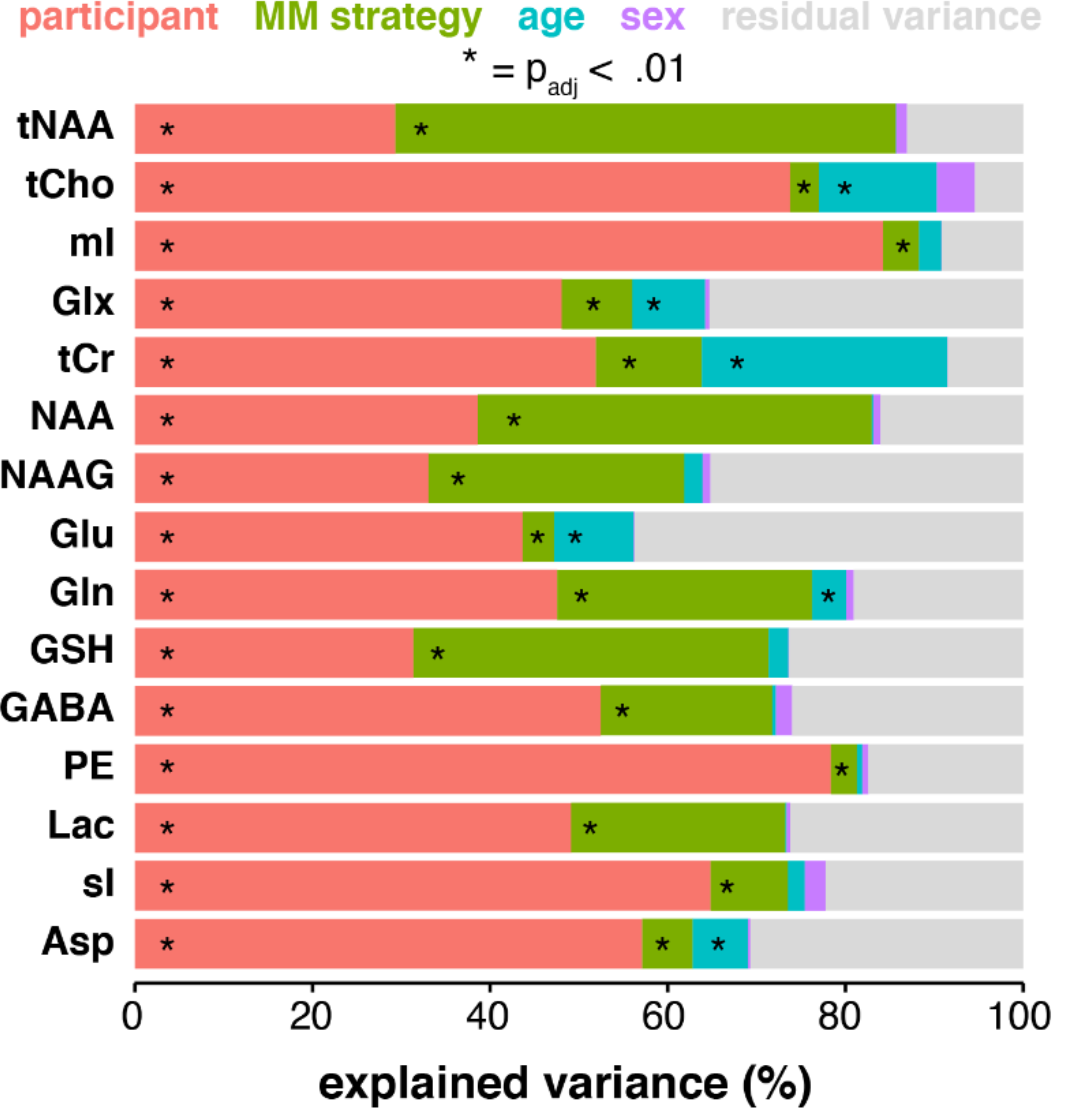
Variance partition analysis for each metabolite across all MM strategies. The linearmixed effect model includes fixed effects for age and sex, and random effects for participant and MM strategy. Significant effects are marked with an asterisk (adjusted p < 0.01 = *).

**Table 2.** Variance partition estimation for each metabolite across all MM strategies. The linear-mixed effect model includes fixed effects for age, and sex and random effects for participant and MM strategy. Significant effects are marked with an asterisk (adjusted p < 0.01 indicated in bold).

| metabolite | participant | MM strategy | age | sex | residual |
| --- | --- | --- | --- | --- | --- |
| tNAA | **29.4** | **56.3** | 0 | 1.2 | 13.1 |
| tCho | **73.8** | **3.2** | **13.3** | 4.3 | 5.4 |
| mI | **84.3** | **4.0** | 2.5 | 0 | 9.2 |
| Glx | **48.1** | **8.0** | **8.2** | 0.5 | 35.3 |
| tCr | **51.9** | **11.9** | **27.6** | 0.1 | 8.5 |
| NAA | **38.6** | **44.4** | 0.2 | 0.8 | 16.1 |
| NAAG | **33.1** | **28.8** | 2.0 | 0.9 | 35.2 |
| Glu | **43.7** | **3.5** | **9.0** | 0.1 | 43.7 |
| Gln | **47.6** | **28.7** | **3.8** | 0.9 | 19 |
| GSH | **31.4** | **40.0** | 2.2 | 0.1 | 26.3 |
| GABA | **52.5** | **19.3** | 0.4 | 1.8 | 26 |
| PE | **78.4** | **2.9** | 0.6 | 0.7 | 17.4 |
| Lac | **49.1** | **24.1** | 0.1 | 0.4 | 26.2 |
| sl | **64.8** | **8.7** | 1.9 | 2.3 | 22.2 |
| Asp | **57.1** | **5.7** | **6.2** | 0.2 | 30.7 |
| **mean** | 52.3 | 19.3 | 5.2 | 0.95 | 22.3 |

## Discussion

We have presented a comparison of six different strategies to incorporate MM basis functions into short-TE LCM metabolite estimation applied to a large lifespan cohort. The major goals were: 1) to compare metabolite level estimates obtained with cohort-mean and subject-specific measured MM and different MM parametrizations; 2) to identify potential sources of differences between the strategies and how they affect metabolite-age associations. The key findings of our study are:

- Group-mean metabolite level estimates significantly differ between parameterized and experimentally derived MM strategies, most notably for tNAA, tCr and GSH. However, for most metabolites, the estimate variance did not change.
- Despite differences in the mean, metabolite estimates correlated strongly (r > 0.6) between the default MM parametrization strategy pMM_default_ and all other MM strategies. Highly J-coupled metabolites showed weaker correlations between MM strategies than singlet signals.
- Experimentally derived MM strategies had better residuals and parsimony metrics than parameterized strategies, indicating greater agreement between data and model. This is especially intriguing since measured MM models have fewer model parameters and therefore fewer degrees of freedom. However, similar performance could possibly be achieved with more accurately parameterized MM basis functions.
- Metabolite-age associations were relatively stable across MM strategies for metabolites with major singlet signals, however, less so for highly J-coupled metabolites.

Variance partition analysis indicated that up to 56.3% (tNAA) of metabolite level variance can be explained by the choice of MM strategy. This highlights the importance of choosing an appropriate MM modeling strategy on validity and interpretation of the results. The variance partition analysis further substantiated the metabolite-age associations for tCho, tCr, Glx, Glu, and Gln. Incorporating the choice of different modeling strategies into a global analysis may increase the robustness of its findings compared to a single correlation analysis.

In this study we took advantage of Osprey’s modularity and its accessible LCM algorithm to create a workflow for directly incorporating subject-specific MM spectra and to introduce separate Gaussian linebroadening parameters for metabolites and MM. These novel modifications highlight a key strength of open-source LCM workflows, since they would be much more challenging to do in closed-source or pre-compiled LCM algorithms. Our results did not demonstrate any substantial advantage conferred by the use of subject-specific MM data in terms of the stability of the age-metabolite association of tCr and tCho, but lead to substantial changes for tNAA, Glu, Gln, and Asp. However, it remains unclear whether this is an artifact introduced by remaining inaccuracies in the MM processing or whether this is an accurate estimation of true biological variability. This suggests that cohort-mean MM basis functions can be used in homogeneous populations and can be time-efficiently reused in similar populations instead of reacquiring for each study. For clinical populations with an underlying pathology affecting MM patterns, it is required to create a cohort-specific MM basis function for this population to achieve more accurate quantification – in addition to the MM basis function for the healthy controls. As a caveat, it is difficult to conclusively establish how many participants-worth of MM data are needed to generate a truly cohort-representative MM spectrum. In the case of highly heterogenous MM spectra within the clinical population, it may even be beneficial to use subject-specific MM. However, it remains unclear how many participants-worth of MM data would constitute a cohort-representative MM spectrum. An interesting strategy would be to acquire a short metabolite-nulled acquisition in every participant, with insufficient SNR to be individually interpretable, but sufficient when averaged across the cohort to generate an appropriate cohort-specific MM spectrum, instead of one long metabolite-nulled acquisition in only a few subjects. While metabolite estimates were significantly different, we did not find substantial differences in the age-metabolite associations when introducing an additional Gaussian linebroadening parameter for MM. Additionally, model residuals and ι1AIC did not improve, indicating no real advantage to this approach. The significant impact of the choice of MM modeling strategy on the short-TE LCM has been shown in several earlier studies^10–12^. Although these studies are either not performed at clinically relevant field strength (≤3T) or their statistical power is limited^10–12^, we found evidence to support a number of their key findings. Confirming previous reports by Schaller et al.^10^ using the LCModel algorithm^6^, we found a significant increase in Glx estimates when using experimentally derived MM, however, in contrast to their results this effect was dominated by increased Gln estimates while Glu remained unaffected. We further substantiated their findings that NAA and tNAA estimates decrease when using measured MMs or an updated MM parametrization.

This is mostly related to the differences in the amplitude of the MM_2.0_ peak which is notably smaller in the pMM_default_ models than in all measured MM spectra. However, it remains unclear whether the large 2-ppm signals in the mMM data represent ‘real’ MM contributions, or are an artifact or unaddressed metabolite signal from the clean-up process. A recently presented method using strongly diffusion-weighted MRS to determine the MM background yielded much smaller MM_2.0_ signals, more distinct MM peaks in the 3.5 to 4 ppm region, and narrower lines in general^44^ compared to the IR method used in our study or previously used saturation-recovery methods^13^. Another potential approach using IR is to acquire a TI-series or 2D-J-spectra at multiple fixed TI in combination with multidimensional modeling that allows separation of metabolite and MM concentration and relaxation parameters. However, multi-dimensional data are more challenging to model^45^. It is unclear which of the approaches is more appropriate and less error-prone to estimate the MM background. Another similar study on the impact of the MM approach in short-TE LCM by Gottschalk et al.^11^ reported comparable changes in metabolite estimates using the QUEST LCM algorithm^46^ with similar directionality of change, e.g. increase in GSH estimates, when comparing modeling approaches with experimentally-derived MM strategies and the QUEST subtraction approach. Confirming findings from that study, we found evidence for a substantial advantage of the experimentally-derived MM strategies over parameterized approaches for the quantification of lactate, which is typically a very small signal on the shoulder of a broad MM peak around 1.3 ppm^11^. Both studies conclude that effects they observe are unlikely to affect the outcome in MRS application studies, however, there is no analysis directly proving this assumption. Both studies featured comparably small samples (11 for Schaller et al., and 20 for Gottschalk et al.) from a narrow age range, which is a substantial limitation to the generalizability of their findings to clinical and age-related studies. In contrast, we studied the impact of the MM strategy choice in a large cohort across the lifespan and unequivocally showed a major influence on the observed metabolite-age associations, especially for highly J-coupled metabolites with strong relevance in aging (e.g. GSH).

A third study by Hofmann et al.^13^ used a saturation-recovery-based method to measure metabolite and MM spectra simultaneously including 40 healthy subjects with three age groups (< 25, 25-55, >55 years), but did not report metabolite quantification results. In comparison to the most widely used LCM algorithm LCModel, Osprey’s algorithm uses an unregularized spline baseline model, no prior distributions for the Lorentzian lineshape parameters, and single Gaussian MM basis functions for the parameterized approach. The advantages of using a single MM basis function may be more pronounced for this less constrained implementation as it strongly reduces the parameter space for the LCM algorithm. This additionally explains the relatively poor performance of the pMM_updated_ strategy, e.g., for the MM_0.91_ peak as the updated strategy only includes a single basis function around 0.9 ppm as opposed to two for the pMM_default_ strategy (MM_0.92_ & Lip_0.9_). Additionally, the original LCModel algorithm includes range of frequencies and FWHM for the Gaussian peaks, as well as expectation values for the Lorentzian lineshape parameters which introduce more freedom to the algorithm to model those signals compared to Osprey. It would be interesting, but beyond the scope of this study, to compare different LCM algorithms, as previously^5^ but with the measured MM spectra. It is possible that the introduction of a regularized baseline such as in LCModel or more advanced baseline approaches as in ABFit^47^ or elsewhere^12^ could improve the performance of parameterized MM estimation during short-TE LCM and allow for separation of metabolite, MM, and baseline signals. Apart from regularization terms the flexibility of a spline baseline can also be controlled by the density of the spline knot spacing and has been shown to have major impact on the quantification^48,49^. This is especially the case if separate Gaussian MM basis functions are used. In this study we used Osprey’s default spbl knotspacing of 0.4 ppm which resulted in moderate baseline flexibility for the experimentally derived MM strategies and a good agreement in the combined [total MM + spline baseline] model for all strategies. However, increasing the spbl knotspacing could potentially further improve the performance of the experimentally derived MM strategies as the underlying MM signals are sufficiently described by the single MM basis function.

MM spectra have much broader, smaller, and less well-defined signals than metabolite spectra, and they inevitably still contain contributions from metabolites that are not perfectly nulled, particularly in a single-inversion recovery experiment because of differences in T_1_ relaxation. Therefore, processing, removal of residual metabolite signals (clean-up), and quantification are crucial, but may suffer from imperfections. Potential issues are accurate frequency referencing due to the broader signals, residual metabolite signals due to imperfect metabolite-nulling^10,11^ and their insufficient removal during post-processing, or an insufficient number of splines to generate the noise-free clean MM spectra. While there were no obvious residual metabolite signals in the MM spline model, the higher variance of tNAA and Glx estimates for the subject-specific mMM strategies suggests greater variability of residual metabolite signals or imperfect clean-up of the single-subject MM spectra. T_1_ differences between moieties may be implicated, and the use of moiety-specific basis functions during clean-up might reduce residual metabolite signals. Double-inversion recovery approaches might improve the consistency of metabolite nulling, however, at the cost of greater T_1_-weighting of the MM signals. To reduce T_1_-weighting effects diffusion-weighted metabolite-nulling could be used which is less susceptible to T_1_ differences between metabolites or introduced by aging^44,50^. Additionally, the MM derived with metabolite nulling will differ from the MM background in the metabolite spectra by a T_1_ relaxation factor. This is accounted for during LCM with a single linear amplitude scaling parameter for the MM basis function. While this is consensus-recommended best practice, it obviously requires the assumption that all MM signals relax with the same T_1_, which may not be justified. Finally, another interesting approach to account for MM signals is to generate MM-nulled spectra with multiple-IR methods^51^. While this approach reduces the overall SNR of the spectra it significantly reduces the contributions of MMs, does not require separate acquisition of MM data, and could therefore improve the quantification of the remaining metabolite signals.

### Perspective for extension of multi-LCM approaches

This study is, to the best of the authors’ knowledge, the first study to explicitly consider the additional variance introduced from different LCM strategies and settings in the final statistical analysis to model biological effects in the cohort, in this case metabolite-age associations. This ‘meta-LCM’ approach could prove to be more robust to model-based bias than a statistical analysis based on a single model, as seen in the differences observed in the separate correlation matrix. A novel generalized LCM analysis fully capturing model-related variance should aggregate a large number of diverse modeling strategies, in contrast to the few drastically different approaches presented here, which resulted in a large portion of the overall variance being attributed to the MM strategy for several metabolites. These diverse strategies may include different signal parametrizations (arbitrary lineshape as in Osprey^14^ and LCModel^6^, asymmetric lineshape as in ABFit^47^, number of lineshape parameters), MM approaches (Gaussian basis functions, Voigt basis functions, mean/individually measured MM spectra), background signal descriptions (spline functions as in Osprey^14^ and LCModel^6^, polynomials like in FSL-MRS, subtraction techniques as in QUEST^15,52^ and Tarquin), parameter regularization, and model domain (time vs frequency domain, combined time and frequency domain, real- or complex-valued data). With ever increasing computational resources, a ‘multi-model’ meta analysis could lead to more robust metabolite estimation and provide a better estimate of estimation uncertainties related to the choice of modeling strategy.

### Limitations

A major limitation of the study, and in-vivo MRS in general, is the absence of a ‘ground truth’ of metabolite level estimates to validate LCM results against. As in our previous studies^5,53^, we evaluated the performance of the different modeling strategies from a combination of estimate variance, fit residuals, parsimony scores, and robustness of observed metabolite-age associations. However, low variance or AICs do not reflect higher accuracy and may rather indicate insufficient responsiveness to real biological variance. The use of simulated spectra as a ground truth is promising as a validation strategy^54^, but can only succeed if synthetic data are truly representative of in-vivo conditions. Another limitation is the single LCM algorithm Osprey that we used here, which, while conceptually and in performance^5^ similar to LCModel, does not feature regularization terms on the baseline or the lineshape, and features fewer metabolite soft constraints. The data in this study are of exceptionally high spectral quality (high SNR, narrow linewidth, and no lipid contamination) and our observations and conclusions may substantially differ for spectra of lower spectral quality (low SNR, broader linewidth, and lipid contamination). Systematic differences in the baseline due different water suppression schemes could affect the model performance. However, such differences are accounted for by the spline baseline and could explain differences in the background signals of the metabolite and the metabolite-nulled spectra. Finally, since this study focuses on MM modeling strategies, we did not include a detailed interpretation and discussion of associations between metabolite levels, age and sex, which can be found elsewhere^9^.

## Conclusion

Inclusion of a single high-SNR MM basis function derived from the cohort-mean of measured metabolite-nulled spectra lead to more robust metabolite estimation than Gaussian parametrizations with more degrees of freedom. Further, it can be included in future studies of healthy subjects making it more time-efficient than single-subject acquisition with minimal impact on metabolite quantification. While there were systematic differences of certain metabolite estimates (tNAA, Glx) between MM strategies, metabolite-age associations (particularly tCho, tCr) were relatively stable. Agreement between MM strategies was generally stronger for singlet signals than for J-coupled signals.

## Abbreviations

LCM: linear-combination modelling
MM: macromolecular
AIC: Akaike information criterion
mMM: measured MM
pMM: parameterized MM
PCC: posterior cingulate cortex
FWHM: full-width half-maximum
HSVD: Hankel singular value decomposition
tMM: total macromolecules
LME: linear mixed-effect model
ICC: Inter-class correlation coefficients
spbl: spline baseline

## Declaration of competing interests

The authors have nothing to declare.

## Acknowledgement

This work is supported by NIH grants R00 AG062230, P41 EB031771 R01 EB016089, R01 EB023963, R21A G060245, and R21 EB033516.

## Notes

https://osf.io/7dxnm/

### Competing Interest Statement

The authors have declared no competing interest.

### Summary of Updates

We have included an additional discussion paragraph to emphasize data homogeneity to be one criterion to successfully use the cohort-mean MM spectra for modelling and how to deal with this aspect in heterogeneous cohorts. We have further included a new analysis to assess differences in the between-MM-strategy correlations and to investigate the stability of the metabolite-age associations. We followed the minor comments to further improve the accessibility of the results and figures.

