## Supplementary Material for "Cohort-mean measured macromolecules lead to more robust linear-combination modeling than parameterized and subject-specific ones"

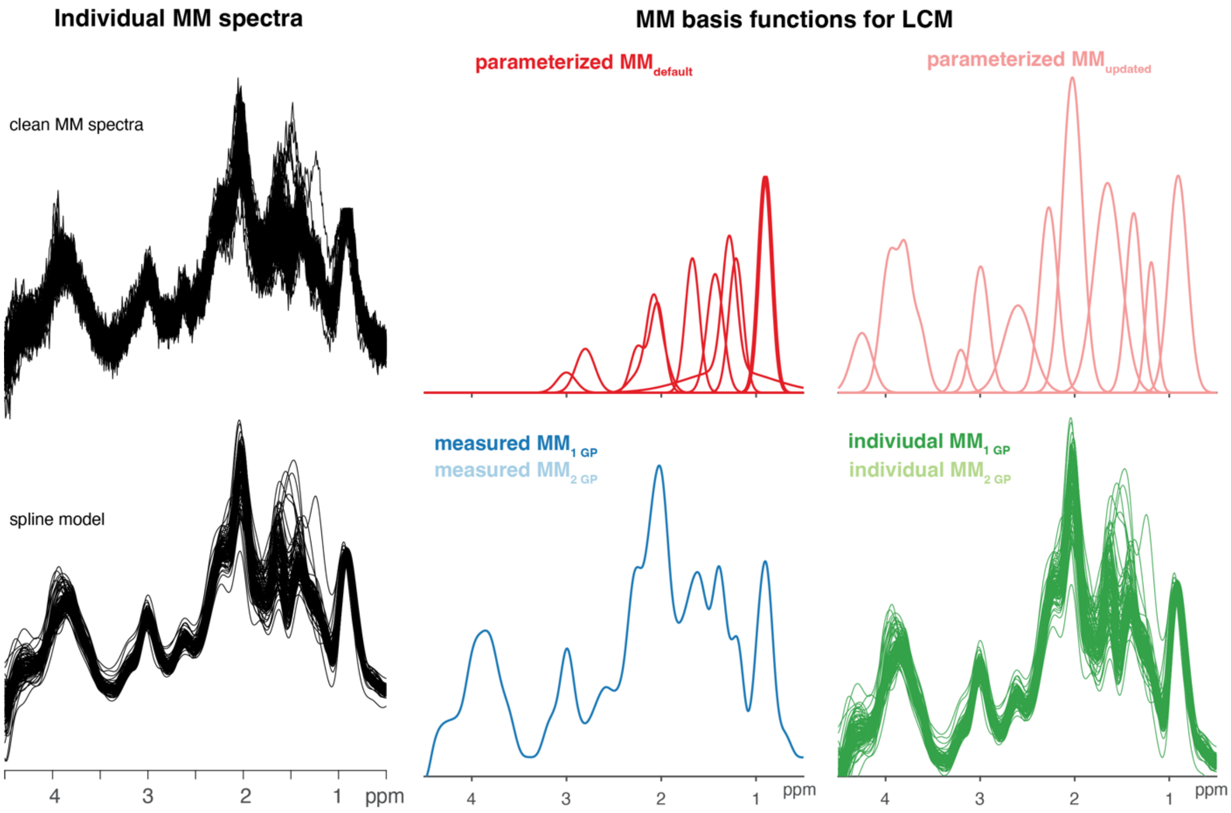


*Supplementary Material 1 - 'Clean' MM spectra and matching spline models from all subjects. Summary of MM basis function for the short-TE LCM including Gaussian parameterization according to the LCModel software (parameterized $MM_{default}$) and updated Gaussian parametrization derived from the mean MM spline model (parameterized $MM_{updated}$). Single experimentally derived MM basis functions described as the mean of the MM spline models (measured $MM_{1\ GP}$ & measured $MM_{2\ GP}$) and individual MM spline models (individual $MM_{1\ GP}$ & individual $MM_{2\ GP}$)*

| pMM$_{default}$ | | | | | pMM$_{updated}$ | | | | |
|---|---|---|---|---|---|---|---|---|---|
| *Name* | *Frequency [ppm]* | *FWHM [ppm]* | *Amplitude* | *Ratio for Soft Constraint* | *Name* | *Frequency [ppm]* | *FWHM [ppm]* | *Amplitude* | *Ratio for Soft Constraint* |
| MM$_{0.92}$ | 0.91 | 0.14 | 3 | Reference | MM$_{0.92}$ | 0.91 | 0.14 | 3 | Reference |
| MM$_{1.20}$ | 1.21 | 0.15 | 2 | 0.3 | MM$_{1.20}$ | 1.19 | 0.08 | 1.81 | 1 |
| MM$_{1.38}$ | 1.43 | 0.17 | 2 | 0.75 | MM$_{1.38}$ | 1.38 | 0.12 | 2.5 | 1 |
| MM$_{1.7}$ | 1.67 | 0.15 | 0.2 | 0.375 | MM$_{1.7}$ | 1.65 | 0.2 | 2.91 | 1 |
| MM$_{2.0}$ | 2.08 | 0.15 | 1.33 | 1.5 | MM$_{2.0}$ | 2.03 | 0.15 | 4.38 | 1 |
| | 2.25 | 0.2 | 0.33 | - | MM$_{2.27}$ | 2.27 | 0.14 | 2.59 | 1 |
| | 1.95 | 0.15 | 0.33 | - | MM$_{2.6}$ | 2.6 | 0.22 | 1.21 | 1 |
| | 3 | 0.2 | 0.4 | - | MM$_{3.0}$ | 2.99 | 0.12 | 1.76 | 1 |
| Lip$_{0.9}$ | 0.89 | 0.14 | 3 | 0.267 | MM$_{3.2}$ | 3.2 | 0.1 | 0.6 | 1 |
| Lip$_{13}$ | 1.28 | 0.15 | 2 | Reference | MM$_{3.7\text{-}4.0}$ | 3.65 | 0.12 | 0.96 | 1 |
| | 1.28 | 0.089 | 2 | - | | 3.79 | 0.09 | 1.34 | 1 |
| Lip$_{2.0}$ | 2.04 | 0.15 | 1.33 | 0.15 | | 3.95 | 0.14 | 1.95 | 1 |
| | 2.25 | 0.15 | 0.67 | - | MM$_{4.2}$ | 4.2 | 0.15 | 0.74 | 1 |
| | 2.8 | 0.2 | 0.87 | - | | | | | |

*Supplementary Material 2 - Properties of the Gaussian functions of the broad macromolecule and lipid resonances included in the pMM$_{default}$ strategy, taken from section 11.7 of the LCModel manual, and the updated parameterization pMM$_{updated}$ derived from the mean MM spectra . The amplitude values are scaled relative to the $CH_3$ singlet of creatine with amplitude 3.*

### Summary following minimum reporting standards in MRS generated in Osprey

See Lin et al. 'Minimum Reporting Standards for in vivo Magnetic Resonance Spectroscopy (MRSinMRS): Experts' consensus recommendations. NMR in Biomedicine. 2021;e4484. doi.org/10.1002/nbm.4448

| **1. Hardware** | |
|---|---|
| a. Field strength [T] | 3 T |
| b. Manufacturer | Philips Healthcare |
| c. Model (software version if available) | Ingenia CX R 6.0.541 |
| d. RF coils: nuclei (transmit/receive), number of channels, type, body part | 1H |
| e. Additional hardware | - |

| **2. Acquisition** | |
|---|---|
| **Short-TE PRESS** | |
| a. Pulse sequence | Philips PRESS |
| b. Volume of interest (VOI) locations | posterior cinculate cortex (PCC) |
| c. Nominal VOI size [$mm^3$] | 30 x 26 x 26 $mm^3$ |
| d. Repetition time (TR), echo time (TE) [ms] | TR 2000 ms, TE 30 ms |
| e. Total number of averages per spectrum<br>i. Number of averaged specra per subspectrum | 96 total averages<br>- |
| f. Additional sequence parameters | F1: 2000 Hz, 2048 points |
| g. Water suppression method | VAPOR |
| h. Shimming method, reference peak, and threshold of acceptance of shim chosen | $2^{nd}$ order pencil beam, water |
| i. Trigger or motion correction | - |
| **Metabolite-nulled PRESS** | |

| **2. Acquisition** | |
|---|---|
| a. Pulse sequence | JHU patch metabolite-nulled PRESS |
| b. Volume of interest (VOI) locations | posterior cinculate cortex (PCC) |
| c. Nominal VOI size [$mm^3$] | 30 x 26 x 26 $mm^3$ |
| d. Repetition time (TR), echo time (TE), inversion time (TI) [ms] | TR 2000 ms, TE 30 ms, TI 650 ms |
| e. Total number of averages per spectrum<br>i. Number of averaged specra per subspectrum | 96 total averages<br>- |
| f. Additional sequence parameters | F1: 2000 Hz, 2048 points |
| g. Water suppression method | CHESS |
| h. Shimming method, reference peak, and threshold of acceptance of shim chosen | $2^{nd}$ order pencil beam, water |
| i. Trigger or motion correction | - |

| **3. Data analysis methods and outputs** | |
|---|---|
| a. Analysis software | Osprey 2.2.0 |
| b. Processing steps deviating from Osprey | None |
| c. Output measure | TissCorrWaterScaled (Gasparovic et al. 2006) |

| **3. Data analysis methods and outputs** | |
|---|---|
| d. Quantification references and assumptions, fitting model assumptions | Basis set list:<br>Asc, Asp, Cr, CrCH2, GABA, GPC, GSH, Gln, Glu, mI, Lac, NAA, NAAG, PCh, PCr, PE, sI, Tau, NAA<br><br>MM basis functions:<br>$pMM_{default}$: $MM_{0.92}$, $MM_{1.20}$, $MM_{1.38}$, $MM_{1.7}$, $MM_{2.0}$, $Lip_{0.9}$, $Lip_{1.3}$, $Lip_{2.0}$<br><br>$pMM_{updated}$: $MM_{0.92}$, $MM_{1.20}$, $MM_{1.38}$, $MM_{1.7}$, $MM_{2.0}$, $MM_{2.27}$, $MM_{2.6}$, $MM_{3.0}$, $MM_{3.2}$, $MM_{3.7-4.0}$, $MM_{4.2}$<br><br>$mMM_{1\ GP}$ & $mMM_{2\ GP}$: cohort mean measured MM<br><br>$imMM_{1\ GP}$ & $imMM_{2\ GP}$: subject-specific measured MM<br><br>Fitting method: Osprey basline knot spacing 0.40 ppm |

| **4. Data quality** | |
|---|---|
| a. SNR (NAA), linewidth (NAA) [Hz] | SNR: 151 +- 23, linewidth 6.68 +- 0.85 Hz |
| b. Data exclusion criteria | visible lipid/ethanol contamination |
| c. Quality measures of postporcessing model fitting (Mean Relative Amplitude Residual) | $pMM_{default}$: 4.93 +- 1.06%<br>$pMM_{updated}$: 6.47 +- 1.59%<br>$mMM_{1\ GP}$: 4.10 +- 1.02%<br>$mMM_{2\ GP}$: 4.29 +- 1.04%<br>$imMM_{1\ GP}$: 4.13 +- 0.99%<br>$imMM_{2\ GP}$: 4.28 +- 1.04% |
| d. Mean spectrum created with OspreyOverview | Figure 2 |

*Supplementary Material 3 – MRSinMRS summary generated in Osprey*

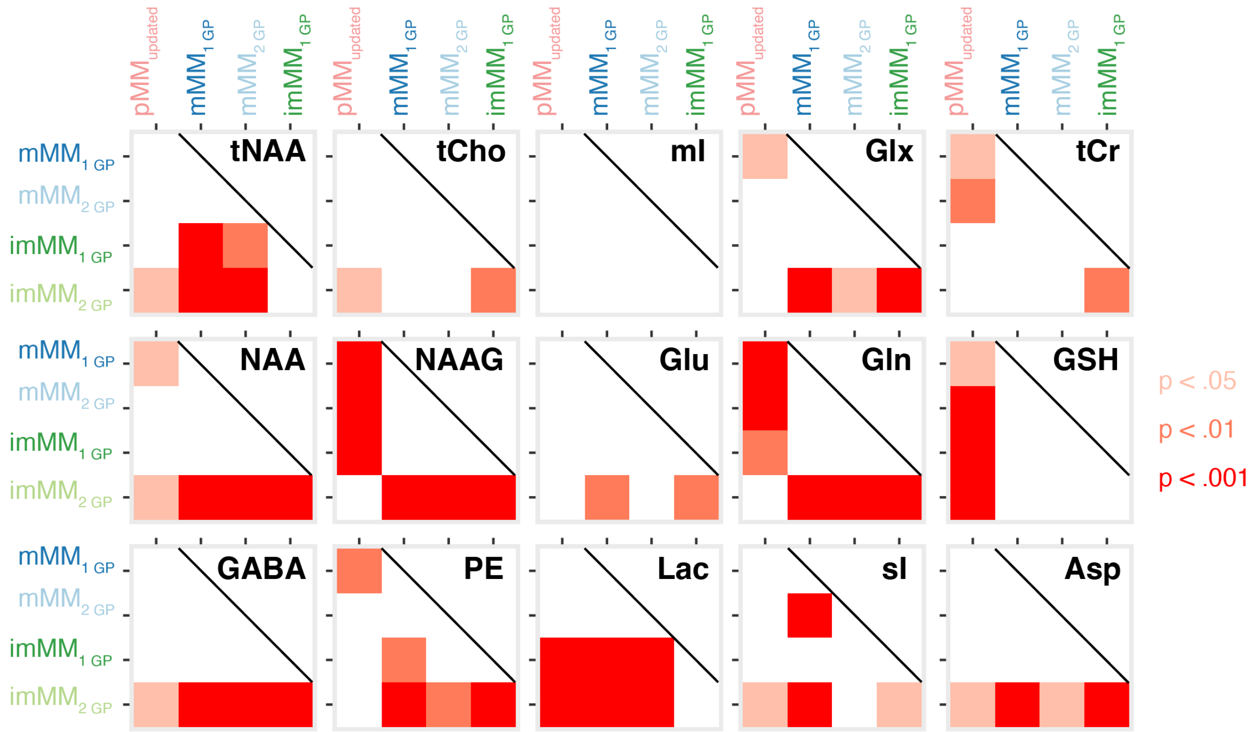


*Supplementary Material 4 – Comparison of the correlation analysis between $pMM_{default}$ and any other MM strategy (see Figure 3C, 4C, and 4F). The Pearson correlation coefficients from the prior correlation analysis were compared for statistical differences for all correlations. Main differences occur for comparisons with $pMM_{default}$ and $imMM_{2\ GP}$.*

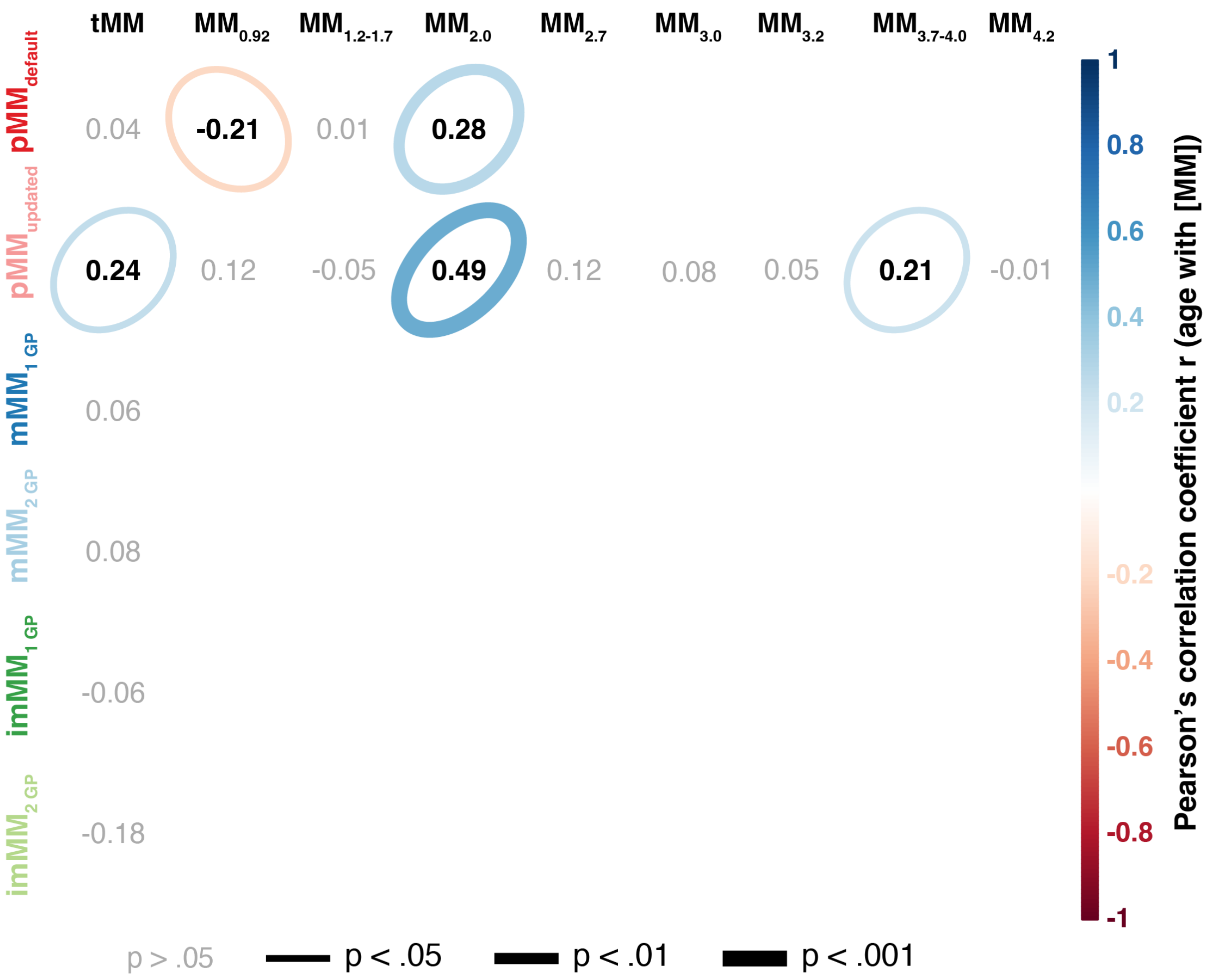


*Supplementary Material 5 - Secondary analysis of MM age associations for all MM strategies. Correlation matrix with Pearson's correlation coefficient r for the metabolite level age association. Significant correlations are indicated as color-coded ellipsoid with the directionality of r indicated by color and orientation and magnitude of r by the size of the ellipsoid. The p values were encoded by the linewidth of the ellipsoid (the smaller the p value the stronger the line). Bonferroni correction is employed for each MM strategy separately*

*(correction factor of 15).*

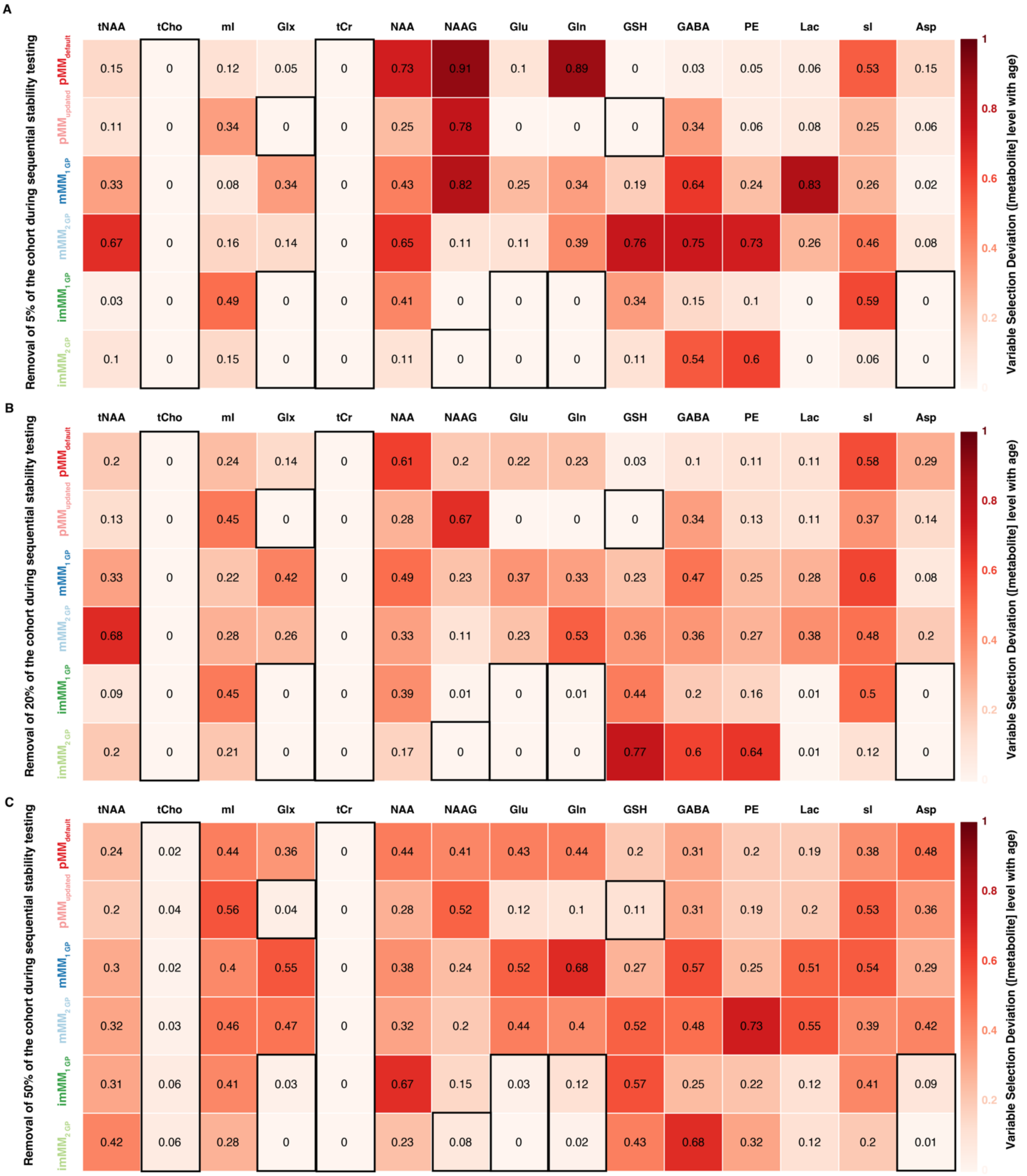


*Supplementary Material 6 – Correlation stability analysis using a variable selection difference (VSD) approach. VSD scores close to 0 indicate a high stablity of the metabolite-age asscoaition. The black frames indicate significant metabolite-age asscoations from the initial correaltion analysis (Figure 6).*
